# Large diversity in the O-chain biosynthetic cluster within populations of Pelagibacterales

**DOI:** 10.1101/2024.03.20.585866

**Authors:** Jose M. Haro-Moreno, Mario López-Pérez, Carmen Molina-Pardines, Francisco Rodriguez-Valera

## Abstract

We have used single-amplified genomes (SAGs) and long-read metagenomics to examine the diversity of the O-chain polysaccharide biosynthesis cluster (OBC) in marine bacteria of the Pelagibacterales order. OBCs are notorious for their diversity and have been used to type strains in pathogens and saprophytes, but their patterns of variation in free-living bacteria are little known. We found that, for these marine heterotrophic bacteria, the diversity is comparable to that of saprophytes, such as Enterobacteriales i.e. nearly each strain carries a different OBC. However, although OBC inheritance was largely vertical, the existence of some shared clusters allowed a comparative analysis. The OBCs diverge along with the genome, which is taken as indicative of old horizontal gene transfer (HGT) events. Only 14 cases of recent HGT were detected and they happened independently of taxonomy or location. Thus, although the O-chain is a major phage receptor in Gram-negative bacteria, the exchange of the complete cluster seems to play a minor role in the phage-bacterium arms-race. By long-read metagenomics, we could detect 380 different OBCs in a single sampling site in the Mediterranean. A single population (single species and sample) of the endemic Ia.3/VII genomospecies had a set of 158 OBCs of which 130 were different. This large diversity in clonal lineages might reflect the large amount of metabolic pathways required to deal with the enormous chemical diversity of dissolved organic matter in the ocean.

## INTRODUCTION

In a recent review (Dittmar et al., 2021) it was calculated that in excess of 400,000 independent metabolic pathways were required to metabolize marine DOM and several hundreds of thousands of ABC transporters have been identified in marine metagenome assemblies (Zhao et al., 2024). Thus, it is to be expected that a single species of a dominant heterotrophic microbe such as Pelagibacter could contain a large diversity of genes in its local pangenome. However, determining the diversity of strains within populations of bacteria has been a major conundrum in microbiology (Viver et al., 2024). Some recent studies in the gut microbiome by culture (Huang et al., 2023), metagenomics (Costea et al., 2017; Sharon et al., 2013; Truong et al., 2017), and long-read amplicons of flagellins (Hu et al., 2022) have concluded that few strains of each species dominate the population within a single individual human microbiome and that they are relatively stable through time. However, studies from metagenomics and single-cell genomics indicated that diversity within a single drop of seawater might be very high (Coleman et al., 2006; Kashtan et al., 2014; Rusch et al., 2007). Nevertheless, even the orders of magnitude of such diversity were hard to establish due to the small size of metagenomic reads or the incompleteness of single-amplified genomes (SAGs). A recent study by culture and metagenomics in an extreme aquatic environment revealed a very high strain diversity in *Salinibacter ruber* with several hundreds of strains within a single pond (Viver et al., 2024).

A very common (if not universal) component of the flexible genome (variable from one strain to another within the same species) is the gene cluster coding for the synthesis of external polysaccharides (Raetz and Whitfield, 2002; Rodriguez-Valera et al., 2016), such as the one containing the genes synthesizing the O-chain of the lipopolysaccharide (often referred to as O-antigen) (Kalynych et al., 2014; Liu et al., 2019; Samuel and Reeves, 2003). They have been known for long in pathogenic/saprophytic bacteria (Huszczynski et al., 2019; Kenyon et al., 2017; Liu et al., 2014; Mostowy and Holt, 2018); their enormous variability has been used to type strains in epidemic outbreaks, i.e. assuming that each strain has a unique combination of exposed polysaccharides. In the order Enterobacteriales, a recent study by Holt and colleagues (Holt et al., 2020) clearly illustrates the patterns of variation found at these organisms’ major glycotype (combinations of exposed polysaccharides) gene clusters. After analyzing more than twenty-seven thousand genomes, they shed light on the evolution of these loci. Their enormous diversity (ca. 18,000 different OBCs were described) and rare exchange between different strains explains their discriminating power used in epidemiology to identify outbreaks produced by the same strain. On the other hand, when HGT was detected the divergence of the genes indicated old transfers (long-term preservation), rare events that do not affect the association between OBC and strain (Holt et al., 2020). These results also discredit the view in which diversity of OBCs is explained by phage-host arms-race that would require fast exchange of the OBCs.

Here we have focused on a study of O-chain biosynthetic gene cluster (OBC) diversity in the marine order Pelagibacterales, taking advantage of the abundance of this oligotrophic microbe in SAG databases and the use of long-read (PacBio HiFi) metagenomics. Their photoheterotrophic lifestyle is well known and makes them among the most relevant microbes in nutrient fluxes in the oligotrophic ocean (Brown et al., 2012; Giovannoni, 2017; Grote et al., 2012; Wilhelm et al., 2007). Besides, their streamlined genome size of ∼1.3 Mb (one of the smallest for planktonic free-living microbes) (Giovannoni et al., 2005; Grote et al., 2012), simplifies the analysis, as they contain a single gene cluster involved in O-chain biosynthesis (e.g. no capsular envelope has ever been found and they have no flagella). Furthermore, the cluster is bounded by the two parts of the ribosomal RNA operon 16S-23S on one side and 5S on the other (Wilhelm et al., 2007), which provides good markers for bioinformatic searches. In addition, a fine taxonomy derived from the ITS can be used to classify the genome with high reliability (García-Martínez and Rodríguez-Valera, 2000) and the rRNA genes can be used as hallmarks for identifying Pelagibacterales reads in long-read metagenomes obtained from the same location in off-shore Mediterranean waters (Haro-Moreno et al., 2021; Zaragoza-Solas et al., 2022). This way we have been able to dissect the diversity of OBC gene clusters as a whole and then also at the population (one single species in one single sample) level. Our findings about the diversity of these OBCs indicate that indeed local strain diversity can be very high even at the single population level.

## RESULTS

### OBCs variation across the order Pelagibacterales

Our first objective has been the analysis of the evolutionary dynamics in the available diversity of Pelagibacterales OBCs, to check if the pattern described previously for saprophytic microbes (Enterobacteriales) could be extrapolated to free-living ones that never interact with immune systems. We screened a dataset comprising nearly 1,700 marine SAGs (Berube et al., 2018; Haro-Moreno et al., 2020; Pachiadaki et al., 2019; Thompson et al., 2019; Thrash et al., 2014) and 20 marine isolate genomes from the whole order Pelagibacterales (clades Ia, Ib, Ic IIa, IIb and IIIa) (**Figure S1**). We avoided clades IV and V, the latter also known as HIMB59, since they have been recently classified as different orders (Haro-Moreno et al., 2020; Molina-Pardines et al., 2023; Viklund et al., 2013). By using the 16S-ITS-23S rRNA operon and the 5S rRNA gene at the left and right ends, respectively, we could recover 806 OBCs >10 Kb long (**Figure S1A**). The Pelagibacterales are characterized by unusually high synonymous replacements (López-Pérez et al., 2020a) and thus the 95% average nucleotide similarity (ANI) used for bacterial species definition (Konstantinidis and Tiedje, 2005) gives a very split taxonomy that contrasts with their high synteny and coverage values (Haro-Moreno et al., 2020). Thus, we have used a genomospecies classification based on phylogenomics and environmental distribution described in (Haro-Moreno et al., 2020).

After manual curation, we found an exception of the location of the OBC in genomes from the genomospecies Ib.4, where there was a genomic rearrangement, that in this clade was found between the 16S-ITS-23S at one end (like in the other clades) but at the other end was located near a cluster of genes involved in the peptidoglycan biosynthesis (**Figure S2A**) and two tRNAs coding for valine and methionine located ∼330 Kb distant from their location in the other clades. A total of 27 SAGs, all in the Ib.4, contained the relocated island. Given that there is no complete genome available for this genomospecies, we collected into a single contig three almost identical (>99 % ANI) SAGs into a partially complete (1.06 Mb) and admittedly chimeric construct (López-Pérez et al., 2020b; Roda-Garcia et al., 2023). **Figure S2B** shows the reconstructed genomic fragment, indicating the position of the 16S, 23S, and 5S rRNA genes. Metagenomic under-recruitment confirmed that the region in **Figure S2A** corresponded indeed to the large OBC found in other clades and that it has been likely translocated to one of the tRNAs, a frequent target of site-directed non-homologous recombination. Still, it was used for the subsequent diversity comparisons and showed very similar behavior (see below).

One hundred and sixty-three islands (one-fifth of the dataset) were categorized as complete, including eight from the rearranged OBCs found in Ib.4. The remaining were partial islands, from which we could recover only the left-hand side (∼23 %), right-hand side (∼22 %), or both sides but in different contigs within the same SAG (∼35 %), (**Figure S1A and Table S1**). Regarding their origin, nearly half of the OBCs (#373) were recovered from a single sample in the Bermuda Atlantic Time-series Study (BATS) (sample SWC-09) collected from a 10 m deep water column sample (**Figure S1B**) (Pachiadaki et al., 2019). Therefore, we can consider all these SAGs as belonging to a single community, and those within a single species, as members of a single population. The remaining OBCs came from a variety of samples from the Atlantic and Pacific Oceans (#170 and #197, respectively) (Berube et al., 2018; Pachiadaki et al., 2019), and from the Mediterranean (Haro-Moreno et al., 2020) (#33) and the Red Seas (#15) (Thompson et al., 2019) (**Figure S1B**). We classified the Pelagibacterales genomes containing OBCs as members of the subclades Ia and Ib (∼90 % of the OBCs), and in much smaller numbers to subgroups Ic, IIa, IIb, and IIIa (**Figure S1C**). This was to be expected since most samples were collected from surface waters, where members of subclades Ia and Ib are largely predominant. Members of subclades Ic and II are mainly found in meso- and bathypelagic waters (Giovannoni, 2017; Thrash et al., 2014; Tsementzi et al., 2016), although some members of the subclade IIa have been detected in surface waters (Bolaños et al., 2022).

The analysis of the 163 complete OBCs showed a large size range, from 10 to 90 Kb (average size 46 Kb) (**Figure S3 and Table S1**). The number of encoded genes varied from 8 to 101, with an average of 45.7 genes per OBC. Grouped by taxa, the average OBC size varied slightly for members of subclades Ia.1 (53 Kb), Ia.3 (51 Kb), and Ia.4 (45 Kb), while subclades Ib.1 (n=13, 57 Kb), and IIa (n=11, 27 Kb) had the largest and smallest glycosylation islands, respectively (**Figure S3A**). OBCs tend to have a GC content lower than the average of the genome in all Gram-negative bacteria (da Silva Filho et al., 2018). For the Pelagibacterales SAGs described here, regardless of the clade considered, the value for the complete OBCs GC content was around 23.8 %, statistically significant (paired t-test, p-value 1e^-116^), lower than the average GC content of the genomes (29.4 %) (**Figure S3B**). The median intergenic spacer (very small in streamlined genomes) was also statistically different between genomes and their OBCs (paired t-test, p-value 0.003), due to the high dispersion of their values (**Figure S3C**). Both differences might reflect the presence of genes of exogenous origin within the OBCs, although it seems difficult to find many microbes with lower GC content than the Pelagibacterales.

### Similarity of shared OBC-types

To evaluate the presence of similar OBC ORFs among the Pelagibacterales genomes, we performed an all-vs-all comparison among them (see methods) (**Table S2**). As shown in **Figure S4**, the majority of the comparisons showed no similarity, measured as the fraction of orthologous genes shared among two OBCs at 50 % AAI and 70 % coverage. For instance, 75.7 % of the OBC pairs shared less than 5 % of orthologs, while this number increases to 94.4 % if we consider those with less than 15 % of orthologous genes **(Figure S4)**. These results illustrate the enormous diversity of OBC genes within the Pelagibacterales order, similar to Enterobacteriales, where 18,384 OBCs resulted in only 2,654 (14.4 %) coding for the same gene sets (Holt et al., 2020). We thus consider two OBCs to belong to the same locus type (hitherto OBC-type) similar to (Holt et al., 2020) if they share at least 90% of the genes. As a result, from the initial set of 806 OBCs (including incomplete ones), 208 genomes shared the OBC-type with another genome in our dataset, while 598 genomes had unique (singleton) OBCs. This is typical of the so-called replacement flexible genomic islands that, although coding for a similar function or (in this case) structure, have completely different gene make-up (López-Pérez et al., 2014; Rodriguez-Valera et al., 2016). An advantage of this kind of genomic island for comparative analysis is that incomplete clusters can be used in pairwise comparisons assuming that similarity throughout some genes can be taken as a sign of a common OBC and likely also a similar (or identical) polysaccharide or glycotype. This is important when working with SAGs, which tend to be fragmented and incomplete, and even more so for long metagenomic reads (see below).

As expected, either the clusters are syntenic and have >85 % similarity or they are completely different in gene content with nearly no orthologous genes detected **(Figure S4)**. For the OBCs belonging to the same OBC-type, in most cases, similarity was very high over the whole stretch, but there were exceptions. For example, **Figure S5** shows in one of the pairs similarity has decreased throughout the right-hand side end of the OBC. In other cases (**Figure S5**) variation seems to be concentrated at the left-hand-side end. In these two cases (and in most) variability seems to increase at the ends (one of them), as was the case in the Enterobacteriales (Holt et al., 2020). This fact improves the reliability of partial sequence comparisons as was done with the metagenomic long reads (see below).

**Figure S6** shows an average amino acid identity (AAI) cladogram tree (Konstantinidis and Tiedje, 2005) rather than the more common average nucleotide identity (ANI). AAI relationships are helpful when comparing highly divergent genome sequences (Barco et al., 2020), as those analyzed here for the whole Pelagibacterales order. **Figure 1A** only shows the genomes sharing OBC-types. A somewhat expected observation was that in most cases SAGs sharing the OBCs were also >99 % AAI overall i.e., they belonged to the same clonal frame or genomovar (Viver et al., 2024). This result was not surprising, since this is the fundament of strain serotyping of Gram-negative bacteria using O-antigens (Kauffmann, 1947; Liu et al., 2019). However, a significant number of associations (∼13 %) were found between individuals belonging to different genomospecies from the same family, or even from different families (**Figure 1A**). There seems to be a bias for certain groups regarding the number of common OBC-types. For instance, genomospecies Ia.3/V (also named as gWID given that has a widespread oceanic distribution by metagenomic fragment recruitment (Delmont et al., 2019; Haro-Moreno et al., 2020)), is one of the genomospecies with the highest number of retrieved genomes (#79, **Figure S6**) but barely showed any shared OBC-type (#7, **Figure 1A**), six of them all corresponded to the same genomovar. Whereas Ib.1, with 87 genomes, had 37 sharing events, five of them jumping across different genomovars. Many shared OBC-types were detected only among members of the same genomospecies, and often between genomes with >95 % AAI (**Figure S7**). Only 49 OBC-types were found across two different genomospecies (31 of the pairs having more than 95 % AAI), while only a single OBC-type was found in four different genomospecies and one in three (**Figure S7)**. Most shared OBCs had the same gene content or only varied in one gene, regardless of the dissimilarity between genomes sharing the OBC (**Figure 1B**). Lastly, the rate of shared OBC-types (**Figure 1C**) drops dramatically below 95 % AAI, regardless of the taxa, and was negligible (24 OBCs) in genomes with less than 90 % AAI. The same pattern applies even when considering different taxa individually i.e. members of the same genomospecies with less than 90 % AAI very seldom share OBC-type.

**Figure 1.**
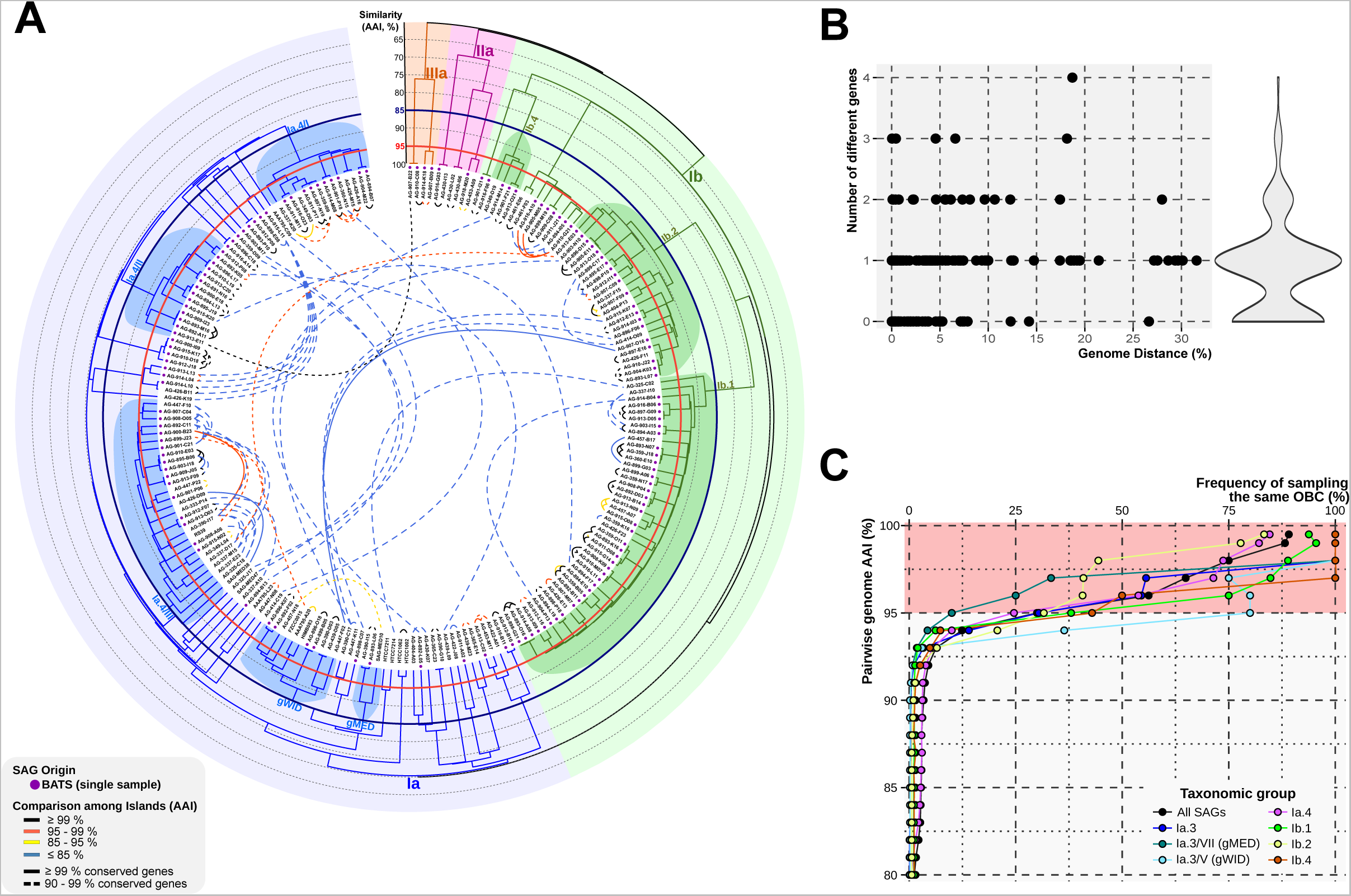
**A.** Cladogram-based classification of the 806 genomes that shared OBC-type, (the complete cladogram is shown in **Figure S6**). The outer rings represent the scale of the cladogram, measured as AAI among genomes (hierarchical clustering, average linkage). Red and blue circles mark the Pelagibacterales limits for species suggested in reference (Konstantinidis and Tiedje, 2005) (85% ANI) and the more common standard 95% respectively (at these levels of similarity AAI and ANI are nearly equal). The most relevant species names are used from reference (Haro-Moreno et al., 2020) and labeled at the corresponding branching line. Inner connections (>90 % shared genes, dashed line, >99 % – continuous line) indicate shared OBC-types between genomes. Connector colors represent the average identity values of the OBC-shared genes. The major Pelagibacterales subclades are shaded: Ia – blue, Ib – green, II – purple, and IIIa – yellow. A purple dot near the genome name indicates that it comes from a BATS single sample. **B.** Variation of the number of different genes for the same OBC-type between pairs of genomes, expressed as the genomic distance (100 - AAI, %). **C.** Frequency of sampling the same OBC-type as a function of the phylogenetic distance between a pair of genomes.

The finding of O-antigen loci shared among distant families suggests horizontal gene transfer (HGT). However, as seen in **Figure 2A**, for most shared OBCs (82 %), the locus genetic distance expressed as ANI (data not shown) or AAI was similar to or slightly higher than the genome distance. Actually, the dN/dS was statistically significantly higher for shared OBC genes (average 0.14 versus 0.1 for the whole genome, paired t-test, p-value <1e^-15^) indicating a slightly higher rate of positive selection (**Figure 2B**). In any case, the values detected for OBC genes indicate that, if they have been exchanged by HGT, it must have been an old event and for the most part they are vertically inherited (long-term preservation) as described for the Enterobacteriales (Holt et al., 2020). The O-chain is a primary target for phages and might be subject to arms-race rapid evolution i.e. change much more rapidly than the rest of the genome to evade phage predation (Letarov, 2023; Mostowy and Holt, 2018). However, the data indicates that this is not achieved by swapping the whole OBC or is very rare, as only fourteen (∼7 % of the total shared OBCs) had an OBC distance that was at least half of the genomic distance, indicative of recent events (**Figure 2A**). These results reveal that rather than rapidly exchanging their OBCs as some scenarios suggest (Mostowy and Holt, 2018; Rostøl and Marraffini, 2019), most OBCs are vertically transmitted over long evolutionary periods. Arms-race is more likely to act at the level of mutation or recombination affecting single genes.

**Figure 2.**
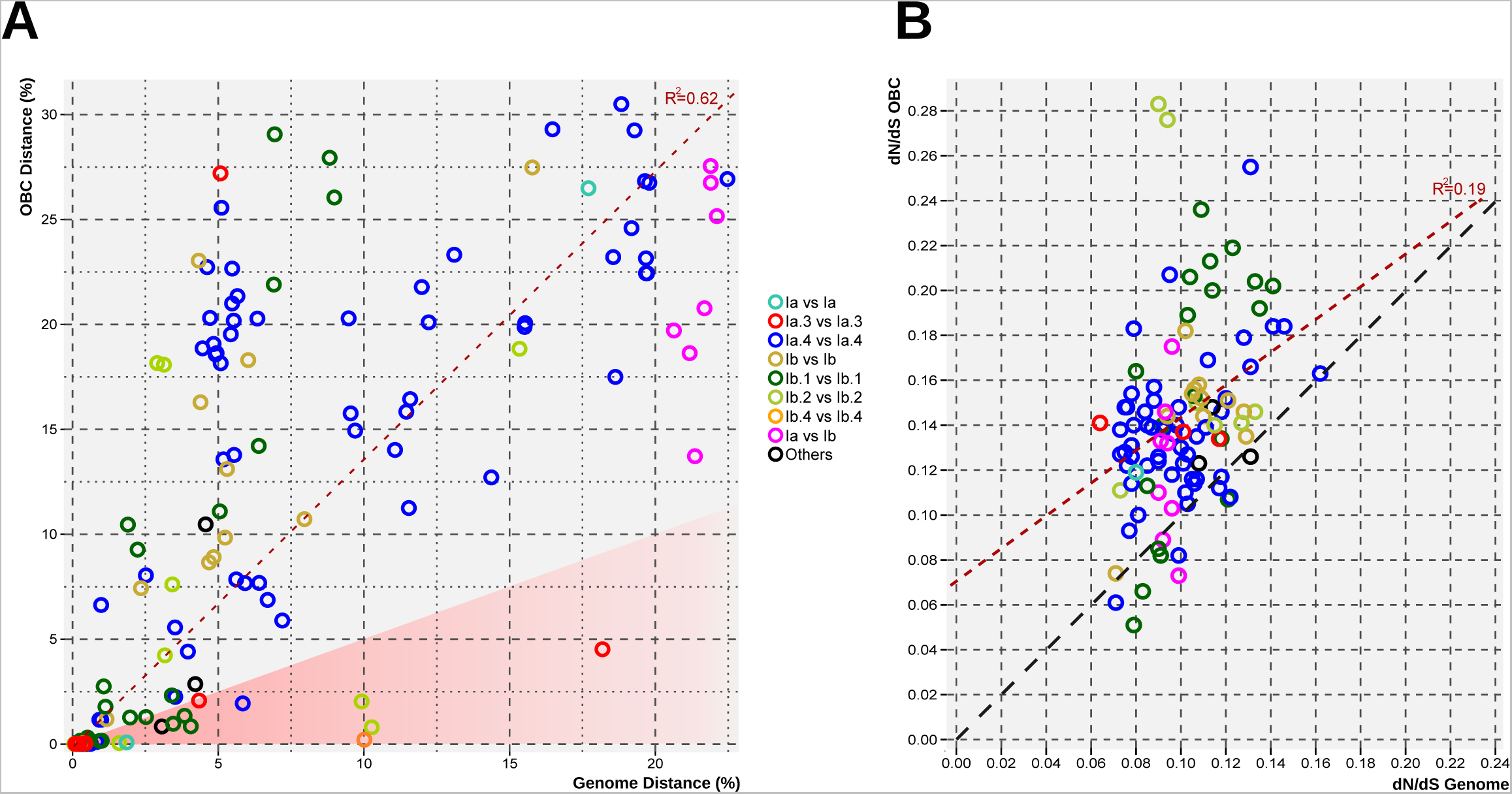
**A.** Scatterplot of the relationship between genome distance and locus distance, both expressed as 100 - AAI (%), for only those genomes sharing an OBC-type. The dashed line in the scatterplot represents the linear regression line. The red-shaded area indicates recent horizontal gene transfer events (genome distance is at least twice the OBC distance). Dots colored by taxonomy. **B.** Scatterplot of the relationship between the dN/dS values for OBCs and genomes. Red dashed line represents the regression line, whereas the black dashed line indicates the y=x.

Given that nearly half of the SAGs in the database came from a single sample of water in the Sargasso Sea (Pachiadaki et al., 2019), we studied how many different OBC-types could be detected at a single location and sample. To do that, we rarefied (Chao et al., 2014; Hsieh et al., 2016) the genomes and the number of different OBC-types detected globally and at BATS. The 373 BATŚ SAGs for the whole Pelagibacterales order had a weak sign of saturation (**Figure 3A**). As expected this coverage was even lower (∼25 %) considering all the 806 genomes for Pelagibacterales as a whole. Therefore, we are far from finding all possible variations in the Pelagibactales OBCs either in the whole ocean or in a single sample. At the genomospecies level and in the single BATS sample, the numbers were too small to allow for a sensible estimation of the total numbers of OBC-types in a single population.

**Figure 3.**
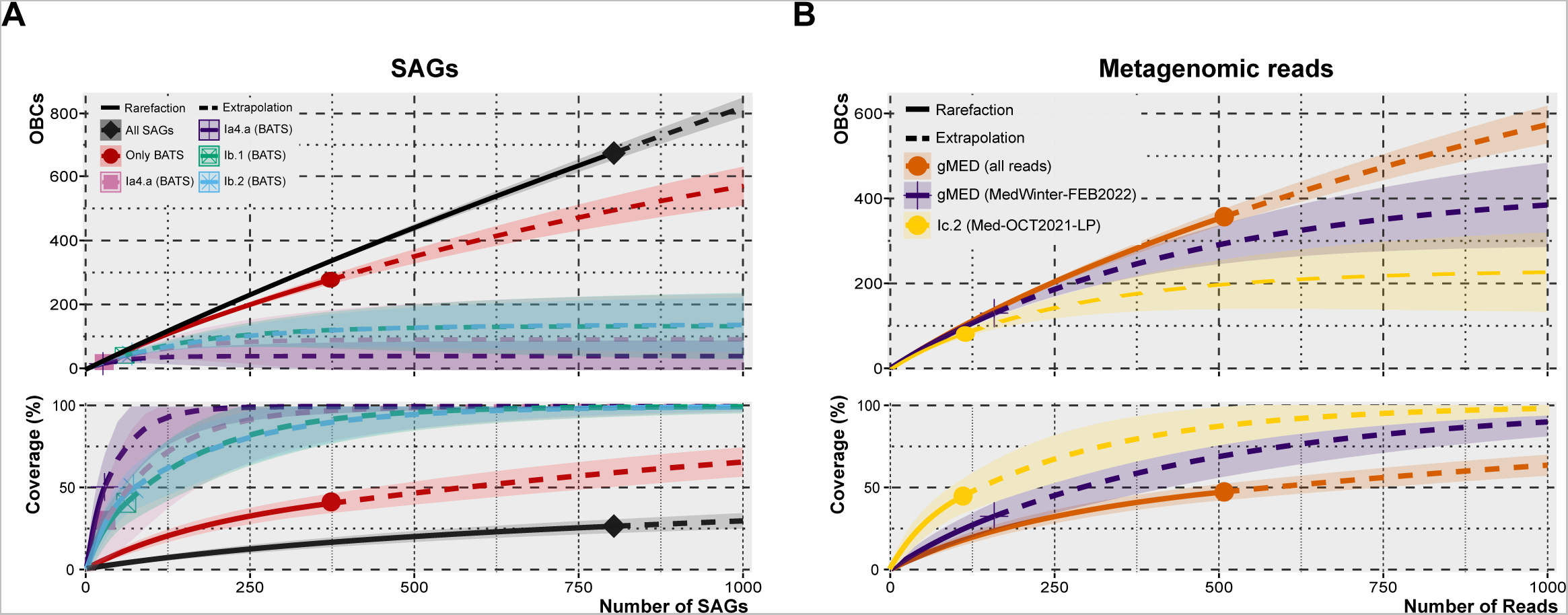
Rarefaction (solid line) and extrapolation (dashed line) curves based on OBC diversity against the number of sequences from **A.** All SAGs (black), and SAGs coming from the single BATS sample, considering all sequences together (red line) or only the four most representative genomospecies (remaining colored lines); **B.** PacBio Sequel II metagenomic reads from a set of Mediterranean samples for two abundant genomospecies, gMED in MedWinter-FEB2022 (purple line), and Ic.2 in Med-OCT2021-75m (yellow line). The bottom plots represent how well the number of detected and predicted sequences covered the diversity of OBCs. The dispersion area shows a 95 % confidence interval.

### Diversity of O-antigen loci within a single population by long-read metagenomics

Metagenomic samples sequenced with PacBio Sequel II can be a good alternative to SAGs to study intrapopulation diversity. Given the large size of the reads, typically between 5 to 15 Kb long (Haro-Moreno et al., 2021), the flexible part of a genome, including fragments presenting part of the OBC can be retrieved before assembly. As discussed before, retrieving only a few genes allows inferring the OBC-type represented by the PacBio read. Thus, we have sequenced and analyzed five different PacBio CCS metagenomes collected from a single location at an off-shore Western Mediterranean Sea off Alicante (see methods). We took samples during winter when the water column is fully mixed (MedWinter-JAN2019 (Haro-Moreno et al., 2021), MedWinter-FEB2022), and during the stratification season at three different layers in the photic zone, the upper photic (Med-OCT2021-15m), the deep chlorophyll maximum (Med-SEP2022-60m, DCM) and the lower photic (Med-OCT2021-75m, below DCM). The prokaryotic community at this location has been thoroughly studied by short and long-read metagenomics before (Haro-Moreno et al., 2021, 2019, 2018; López-Pérez et al., 2017) and the three depths selected when the water column is stratified represent different epipelagic assemblages, as the prokaryotic community significantly differed at these depths (Haro-Moreno et al., 2018). Then, we selected all the reads whose 23S rRNA affiliated within the order Pelagibacterales, and hence the left-hand end of the OBCs could be analyzed to study the variability of the OBC-types within these samples. Only PacBio CCS reads with at least 5 Kb of OBC were used, resulting in a total of 2,780 OBCs from the five metagenomic samples, and the threshold of at least 90 % of orthologous genes was applied for their classification into the same OBC-type. Hence, we found 2,440 OBC-types in all the samples combined (**Figure S8**). The resulting rarefaction, extrapolating to 10,000 reads, showed several OBC-types, between 4,500 and 6,500, for the whole Pelagibacterales order at this single sampling site and over the course of a couple of years with the corresponding seasonal variation (**Figure S8**). This number is certainly in the order of the one found for the order Enterobacteriales (15,730 in 27,000 genomes) (Holt et al., 2020).

To be able to classify the recovered OBCs into genomospecies, we have used the internal transcribed spacer (ITS) between the 16S rRNA-23S rRNA genes. Ecotypes of Pelagibacterales and *Prochlorococcus* have been reliably classified based on their sequence for a long time (Brown and Fuhrman, 2005; García-Martínez and Rodríguez-Valera, 2000; Ngugi and Stingl, 2012; Rocap et al., 2002; Shibl et al., 2014). More recently, a clear association between genomospecies classified by phylogenomics and the ITS cladogram was confirmed (Haro-Moreno et al., 2020). By examining the correspondence between ITS and 23S rRNA gene sequences, we also determined genomospecies from the latter. Therefore, the long-reads with complete ITS (n=1,501) and 23S rRNA sequences (n=2,780) were used to analyze the OBC diversity in single genomospecies (**Table 1**). As expected, the diversity found within the Pelagibacterales CCS reads was astounding, with many sequences grouping with the main phylogenetic groups (Ia, Ib, Ic, and IIa). In the case of the most abundant genomospecies (by number of OBC reads), Ia.3/VII and Ic.2 were detected at significant values (>5 %). Ia.3/VII had 18.4 % of the recovered OBC reads. This group has been characterized as dominating (highest metagenomic recruitment) in epipelagic Mediterranean waters and hence it was also named as gMED (Haro-Moreno et al., 2020). By using only the sequences from a single sample (mixed water column, MedWinter-FEB2022) that had the largest number of OBC reads (158 gMED-affiliated sequences), 130 different OBCs were retrieved in this population, and the rarefaction curve suggested the presence of around 400 different OBC types (294 - 480, 95 % CI) (**Figure 3B**). A similar approach was also applied to another genomospecies, Ic.2 (deep epipelagic), which was found abundant (112 out of 474 OBC identified reads) in the lower photic (75 m deep) sample. The rarefaction curve extrapolates to 254 different OBCs (140 - 310, 95 % CI). Although the numbers are smaller than the ones found for the species *E. coli* as a whole (#800 in (Holt et al., 2020)), the *E. coli* OBCs, although coming from a single well-defined species, were collected from several different populations (e.g. hosts or environments) and the genomic sampling was much larger (ca. 10,000 complete genomes).

**Table 1.**
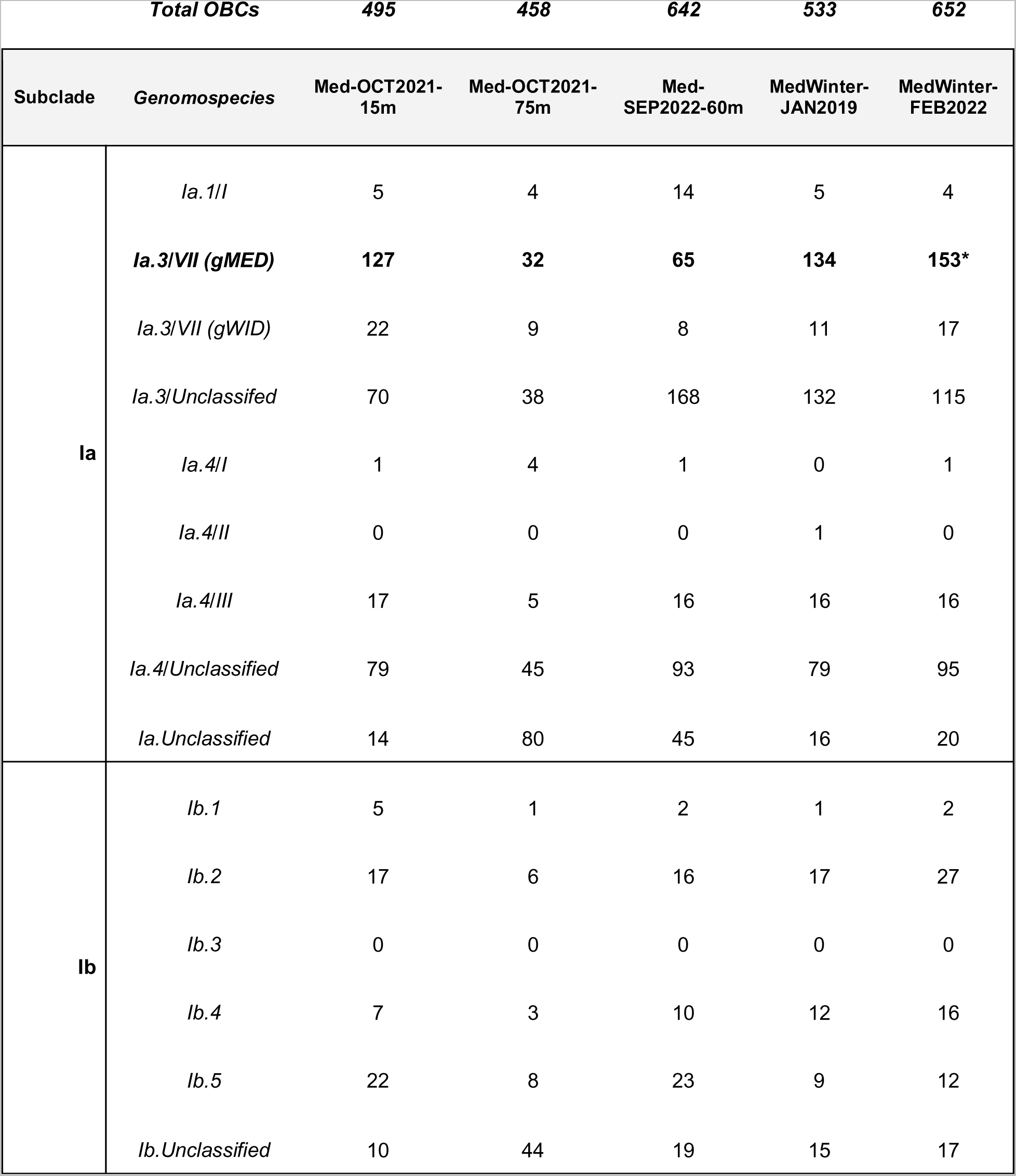
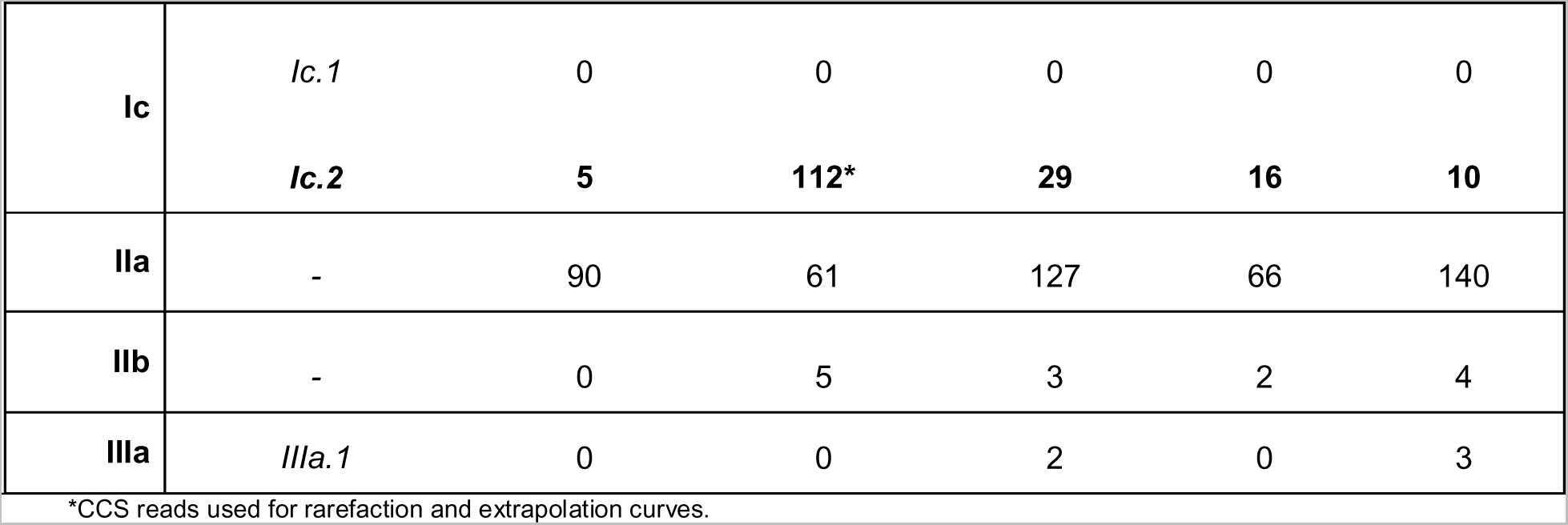
Taxonomic classification of PacBio CCS reads containing a 23S rRNA gene (hence an OBC) and classified to the order Pelagibacterales.

| *Total OBCs* |  | **495** | **458** | **642** | **533** | **652** |
| --- | --- | --- | --- | --- | --- | --- |
| Subclade | *Genomospecies* | Med-OCT2021-15m | Med-OCT2021-75m | Med-SEP2022-60m | MedWinter-JAN2019 | MedWinter-FEB2022 |
| **la** | *la.1/I* | 5 | 4 | 14 | 5 | 4 |
|  | ***la.3/VII (gMED)*** | **127** | **32** | **65** | **134** | **153\*** |
|  | *la.3/VII (gWID)* | 22 | 9 | 8 | 11 | 17 |
|  | *la.3/Unclassified* | 70 | 38 | 168 | 132 | 115 |
|  | *la.4/I* | 1 | 4 | 1 | 0 | 1 |
|  | *la.4/II* | 0 | 0 | 0 | 1 | 0 |
|  | *la.4/III* | 17 | 5 | 16 | 16 | 16 |
|  | *la.4/Unclassified* | 79 | 45 | 93 | 79 | 95 |
|  | *la.Unclassified* | 14 | 80 | 45 | 16 | 20 |
| **lb** | *lb.1* | 5 | 1 | 2 | 1 | 2 |
|  | *lb.2* | 17 | 6 | 16 | 17 | 27 |
|  | *lb.3* | 0 | 0 | 0 | 0 | 0 |
|  | *lb.4* | 7 | 3 | 10 | 12 | 16 |
|  | *lb.5* | 22 | 8 | 23 | 9 | 12 |
|  | *lb.Unclassified* | 10 | 44 | 19 | 15 | 17 |
| **Ic** | *Ic.1* | 0 | 0 | 0 | 0 | 0 |
|  | ***Ic.2*** | **5** | **112\*** | **29** | **16** | **10** |
| **IIa** | - | 90 | 61 | 127 | 66 | 140 |
| **IIb** | - | 0 | 5 | 3 | 2 | 4 |
| **IIIa** | *IIIa.1* | 0 | 0 | 2 | 0 | 3 |

\*CCS reads used for rarefaction and extrapolation curves.

## DISCUSSION

Polysaccharides are extremely versatile macromolecules in which an infinity of structures and properties can be obtained by small modifications in their sugars, bonding order, branching and length. In this sense, they are like proteins but they have to be synthesized by laborious enzymatic steps, exported to the outside of the cell and then polymerized. We have not analyzed the functionality of the genes detected, among other reasons because they lack in most cases reliable annotations beyond the general description of function. But the whole subject of polysaccharide biosynthesis even in model organisms requires more extensive work. Structural predictions and comparisons could help to solve the conundrum but are beyond the scope of this work.

The main objective of this work has been estimating the numbers of strains, i.e. lineages of the same species but with different flexible genomes, present in a single population (same sample). To that end, we have studied the diversity of OBCs within the Pelagibacterales order, as the variation of this region have been traditionally used to type strains (Huszczynski et al., 2019; Kenyon et al., 2017; Liu et al., 2014; Mostowy and Holt, 2018). Unfortunately, we are far from answering this question with the kind of data used here, but we can infer (Hu et al., 2022) that it should be slightly less than the numbers of different OBCs, i.e. in the order of hundreds of strains. This number fits well with the numbers found for *Prochlorococcus* by Kashtan and collaborators (Kashtan et al., 2014) by comparing whole SAGs, and it is again surprisingly high for individuals (cells) belonging to the same species and inhabiting a relatively homogeneous environment. They would certainly allow for dealing with large numbers of substrates and conditions by one single species. It has been estimated that dissolved organic matter (DOM) in the ocean contains in the order of 100,000 different chemical formulas half of which have a half-life time of less than two weeks (Zark et al., 2017). Therefore, the number of genes (e.g. transporters, degradative enzymes) required to cope with such chemical diversity is expected to be very large (Dittmar et al., 2021). Thus, it is not surprising that major consumers of DOM such as Pelagibacterales species have proportionally large local gene pools or pangenomes.

The overall diversity and evolution of OBCs described here in a free-living marine proteobacterial order is very similar to that described using cultures of a heterogeneous group of saprophytic and free-living bacteria like the Enterobacteriales. This proves beyond doubt that the classical view (Lerouge and Vanderleyden, 2002; Reeves, 1995) that O-chain diversity is due to the need of the microbes to change their antigenic specificity as protection from host immune systems is unlikely, even if this is partially true under some special circumstances. Furthermore, the term “serotype” should be replaced by “glycotype” which is more realistic (López-Pérez and Rodriguez-Valera, 2016). The variation at the level of exposed polysaccharides in microbes belonging to the same population has been revealed by metagenomics, culture, and other approaches as a constant feature, at least, for aquatic microbes (Layoun et al., 2024; Roda-Garcia et al., 2023; Rodriguez-Valera et al., 2016), including Gram positives (López-Pérez et al., 2020b; Neuenschwander et al., 2018) and even Archaea (Martin-Cuadrado et al., 2015).

The reasons for such extreme flexibility in structures often critical to the survival of cells in nature are still not clear, but in free-living prokaryotes, the most obvious reason to have a high diversity of these markers, within a single population, is avoid predation by protists or phages. Although some O-chains have been considered more refractory to protist predation than others (Sintes and del Giorgio, 2014), there is little doubt that for tiny cells like those of the Pelagibacterales, the predatory pressure of phages is likely more important. Something along the same lines has been described for the most abundant picocyanobacterium *Prochlorococcus*, and its phages (Avrani et al., 2011; Coleman et al., 2006; Schwartz and Lindell, 2017). The O-chain is a major target for phage receptor-binding proteins and thus there are two potential phage-related explanations for its diversity-arms-race and density-dependent negative selection (Abedon, 2022; Rodriguez-Valera et al., 2009). In the latter, increased predation on abundant OBC-types, maintains a large diversity of receptors, distributing the predation pressure among many. The swapping of O-chains is too slow to be effective in an arms-race scenario and, over long timeframes, seems an unlikely explanation. However, at a shorter timeframe, it is clear that mutation or individual gene or cassette swapping could have a role in the phage-host interaction as seen in several laboratory experiments.

What is the relevance of arms race processes then? In the Enterobacteriales, the gain or loss of a small number of genes (1-3) has been described within relatively short (epidemiological) timescales (Holt et al., 2020). Experimental studies have shown that isolated SNPs can make a strain resistant to one phage and also very small variations in the phage receptor-binding protein can revert the resistance (Schwartz and Lindell, 2017). However, it has been shown also by experimental work in cyanophages that a decrease in sensitivity to one phage can increase the sensitivity to another (Schwartz and Lindell, 2017). Furthermore, significant changes in the O-chain structure or composition in a Gram-negative bacterium can alter their antibiotic sensitivity (Pernitzsch et al., 2021) and also, likely, its nutritional preferences. O-chain mutations are notoriously pleiotropic, as they have multiple phenotypic effects (Martínez de Tejada et al., 1995; Pagnout et al., 2019). In addition, the synthesis of an external polysaccharide requires many steps that are connected (synthesis of sugars, transport and linking to the growing external polymer. All these steps must be coordinated and thus submitted to a complexity limitation to change (Perlovsky and Frank-Kamenetskii, 2002). Finally, we would like to suggest that once reached an equilibrium, any change could lead to a major disruption of the species’ adaptation to its niche. In the evolutionary landscape analogy (Gokhale et al., 2009), it would imply falling from the mountain peak to the valley and is likely to be very infrequent.

## METHODS

### Recovery of Pelagibacterales genomes

To evaluate the presence and the number of glycosylation islands from the whole Pelagibacterales order (NCBI taxonomy ID 54526), a compendium of nearly 4,100 genome assemblies was downloaded from the NCBI database. Prior to the analysis, due to the incomplete nature of MAGs, only assemblies from SAGs and isolates were considered, and their degree of completeness and contamination of SAGs were estimated using CheckM v1.1.2 (Parks et al., 2015). SAGs with > 50% completeness and < 5% contamination were kept. A fast taxonomic classification of genomes was performed using the GTDB-Tk v2.1.0 tool (Chaumeil et al., 2019) using the Genome Taxonomy Database (GTDB) release R207 (Parks et al., 2018). Genomes belonging to clades IV and V (HIMB59) or misclassified were removed from the dataset.

### Phylogenomic classification

Using Phylophlan (Segata et al., 2013), a total of 104 genes (26,134 amino acid positions) were used to classify the 806 Pelagibacterales genomes phylogenomically. Genomes of the HIMB59 order and Rickettsia *spp*. were used as an outgroup. The resulting tree was analyzed using iTOL (Letunic and Bork, 2016). Following the well-established SAR11 nomenclature within subclades, phylotypes, and genomospecies described in (Haro-Moreno et al., 2020; López-Pérez et al., 2020a), we used the median distance between nodes and cophenetic correlation coefficient (interval comprised between 0 and 2) to define them.

### Genome annotation and retrieval of the O-chain biosynthetic gene cluster (OBC)

Prodigal v2.6.3 (Hyatt et al., 2010) was used to predict genes from contigs retrieved from the individual genomes in the curated Pelagibacterales dataset containing 1,700 genomes. Predicted protein-encoded genes were taxonomically and functionally annotated against the NCBI NR database using DIAMOND 0.9.15 (Buchfink et al., 2015) and against COG (Tatusov et al., 2001) and TIGRFAM (Haft et al., 2001) using HMMscan v3.3 (Eddy, 2011). tRNA and rRNA genes were predicted using tRNAscan-SE v2.0.5 (Lowe and Eddy, 1996) and barrnap v0.92 (https://github.com/tseemann/barrnap), respectively.

Contigs containing a signal for the 23S and 5S rRNA genes were selected as they are characterized as markers for the start and ending points of the OBC in Pelagibacterales. If both markers were found within the same contig the island was classified as complete.

### Genomic pairwise comparison

Average nucleotide and amino acid identities (ANI and AAI) between a pair of genomes were calculated using the JSpecies with default parameters (Richter and Rossello-Mora, 2009) and CompareM (https://github.com/donovan-h-parks/CompareM) software packages respectively. A cladogram of the genomes containing the OBC (complete or partial) was constructed using an all-vs-all matrix of the AAI values among genomes, followed by hierarchical clustering with the average Euclidean distance among values with the hclust function in R (R Development Core Team and Team, 2011).

### Determination of O-chain biosynthesis locus types

AAI values between pairs of OBCs were calculated with CompareM, considering 50 % amino acid identity as the threshold to establish similarity among genes. We only considered the pairwise combinations of A (complete OBC) vs A, A vs B (partial OBC), B vs A, A vs C (only left-hand side OBC), C vs A, A vs D (only right-hand side OBC), D vs A, B vs C, C vs B, B vs D, D vs B, C vs C, and D vs D. Two or more OBCs belonged to the same OBC-type if they shared at least 90 % of the genes (considering the smallest OBC).

### dN/dS values from OBCs and genomes

Estimation of the numbers of synonymous (dS), non-synonymous (dN) mutations, and the dN/dS ratio was performed using orthologr (Drost et al., 2015) against the set of genomes sharing an OBC and their OBCs. The OBC cluster was extracted from the genome sequences before calculation. Ortholog genes were detected using a reciprocal best-hit approach using DIAMOND and the dN/dS ratios were estimated using the Comeron algorithm (Comeron, 1995).

### PacBio CCS15 metagenomic reads

Four marine samples were collected from the same sampling site in the epipelagic Mediterranean Sea at 20 nautical miles off the coast of Alicante (Spain) (37.35361°N, 0.286194°W). MedWinter-FEB2022 (20 m deep) was collected during winter, when the water column is fully mixed. Med-OCT2021-15m, Med-SEP2022-60m and Med-OCT2021-75m were collected in summer, during a strong stratification period. We added to the comparison a winter sample collected in January 2019 and sequenced with PacBio Sequel II (MedWinter-JAN2019 (Haro-Moreno et al., 2021)). For each depth, 200 L were collected and filtered on board as described in the study of Haro-Moreno et al. (2018). Briefly, seawater samples were sequentially filtered through 20-, 5-, and 0.22-μm pore filter polycarbonate filters (Millipore). Water was directly pumped onto the series of filters to minimize the bottle effect. Filters were immediately frozen on dry ice and stored at −80°C until processing.

DNA extraction was performed from the 0.22-μm filter (free-living bacteria) following the MagAttract Purification Kit protocol (QUIAGEN). Metagenomes were sequenced using PacBio Sequel II (one 8M SMRT Cell Run, 30-h movie) (Novogene, South Korea). To improve the quality of the PacBio raw reads, we generated Highly Accurate Single-Molecule Consensus Reads (CCS reads) using the CCS v6 program of the SMRT-link package. The minimum number of full-length subreads required to generate a CCS read was set to 15 (> 99.99 % base call accuracy).

### ITS and 23S phylogenies of Pelagibacterales genomes and PacBio CCS reads

PacBio CCS15 reads >5 Kb were screened to detect 16S and 23S rRNA genes using barrnap. Using SILVA (Quast et al., 2013), sequences affiliated with Pelagibacterales were kept and complete internal transcribed spacers (ITS) and 23S rRNA genes were extracted for further analysis. Phylotype classifications based on the ITS and 23S rRNA gene were inferred using the neighbour-joining approach in MEGA11 (Tamura et al., 2021), with 1000 bootstraps and the Jukes-Cantor model of substitution. Phylotype assignment followed existing ITS and 23S nomenclatures (Brown et al., 2012; García-Martínez and Rodríguez-Valera, 2000; Ngugi and Stingl, 2012).

### Rarefaction curves among O-antigen loci

We quantified the number of locus types, aka locus diversity, by applying an ecological modelling approach using iNEXT package in R (Hsieh et al., 2016). In this approach, each SAG or CCS read was considered ecological “sites” and the OBC-types were considered the observed “species” in those sites. The rarefaction curves were calculated by extrapolating our data to 1,000 SAGs and CCS reads for measuring at the genomospecies level, and to 10,000 CCS reads for the whole Pelagibacterales order.

## ACKNOWLEDGEMENTS

This work was supported by grant “FLEX3GEN” PID2020-118052GB-I00 (cofounded with FEDER funds) from the Spanish Ministerio de Economía, Industria y Competitividad to F.R-V and M.L.-P. J.M.H.-M. was supported with a PhD fellowship from Margarita Salas program, cofounded by the Spanish Ministerio de Universidades and the European Union—Next-Generation EU (2021/PER/00020). C.M.-P. was supported by a Ph.D. fellowship from the Spanish Ministerio de Ciencia e Innovación (PRE2021-098122). We are grateful to Ramunas Stepanauskas for suggestions related to SAGs and comments on a draft of the manuscript.

## AUTHOR CONTRIBUTIONS

F.R-V conceived the study. J.M.H-M, C.M-P, and M.L-P collected and processed the metagenomic samples. J.M.H-M, M.L-P, and F.R-V analyzed the data. J.M.H-M and F.R-V contributed to write the manuscript.

## DATA AVAILABILITY

Metagenomic datasets have been submitted to NCBI SRA and are available under BioProject accession number PRJNA1088973 (PacBio CCS15 reads: MedWinter-FEB2022-CCS [SAMN40517308], Med-OCT2021-15m-CCS [SAMN40517305], Med-SEP2022-60m-CCS [SAMN40517307] and Med-OCT2021-75m-CCS [SAMN40517306]).

## SUPPORTING INFORMATION

**Figure S1. A.** Upper pie chart indicates the total number of Pelagibacterales genomes screened (pale color), and the number of genomes on which we could identify the O-chain biosynthetic gene cluster (OBC) (black area). The bottom pie chart distributes the 806 OBCs according to their completeness: A – complete OBC; B – the boundaries of the OBC, i.e. the 23S rRNA gene on the left-hand side and the 5S rRNA gene on the right-hand side, were detected, but in two different contigs from the same genome; C – only the left-hand side; D – only the right-hand side. **B.** Number of OBCs recovered by oceanic region. **C.** Taxonomic classification of Pelagibacterales-containing OBCs based on a maximum-likelihood phylogenetic tree from shared proteins (see methods). The resulting phylogenetic groups follow the nomenclature described in Haro-Moreno et. al., 2020 (Haro-Moreno et al., 2020).

**Figure S2. A.** Genomic comparison of four selected complete OBCs from members of the Ib.4 phylogroup. Note that in this case, the glycosylation island is located between the 16S-ITS-23S rRNA genes and the tRNA-Val, tRNA-Met genes, and the core genes involved in the peptidoglycan biosynthesis. **B.** Reconstruction of a partial genome, Ib4-rB, in a single contig after the co-assembly of 3 nearly identical (>99 % ANI) SAGs. The locations of the *dna*A, 16S, 23S, and 5S rRNA genes are indicated. The metagenomic fragment recruitment of this genome in the Mediterranean Sea (Med-SEP2014-60m) confirmed the location of the glycosylation island.

**Figure S3.** Genomic properties of the 163 complete OBCs and their corresponding genomes. Boxplot on the left summarises the median, first and third quartiles of the length of the OBC, considering all sequences as a unit (leftmost boxplot) or divided by subgroups. Numbers below the taxonomic name indicate the number of OBCs for each group. Boxplots on the right show the difference in the GC content and the intergenic spacer (upper and lower panels, respectively) between the OBC and its genome. Stars indicate the p-value (*** p-value < 0.001, ** p-value < 0.01).

**Figure S4. A.** Histogram resulted from the all-vs-all comparison of the 806 OBCs at 50 % amino acid identity threshold. X-axis indicates the percentage of orthologous genes (OF) of the shortest sequence (genes shared between two OBCs), in groups of 5 % OF.

**Figure S5.** Examples of OBC-type sharing among Pelagibacterales genomes. OBCs are aligned and AAI color-coded.

**Figure S6.** Cladogram-based classification of the 806 genomes containing an OBC, represented as in **Figure 1**.

**Figure S7.** Number of OBCs found in one or several taxonomic groups (>85 % AAI or >95% AAI).

**Figure S8.** Rarefaction (solid line) curves based on OBC diversity against the number of sequences from a set of five PacBio Sequel II metagenomic reads collected from the Mediterranean Sea.

**Table S1.** Summary of Pelagibacterales genomes (SAGs and isolates) with an O-chain Biosynthetic gene Cluster (OBC).

**Table S2.** Summary of pairwise comparisons among O-chain Biosynthetic gene clusters (OBCs).

