## Supplementary Figures for "Large diversity in the O-chain biosynthetic cluster within populations of Pelagibacterales"

[1]Evolutionary Genomics Group, División de Microbiología, Universidad Miguel Hernández, Apartado 18, San Juan 03550, Alicante, Spain.

Universidad Miguel Hernández, División de Microbiología, Apartado 18, San Juan de Alicante, 03550 Alicante, Spain.

**This PDF file includes:**

Figures S1 to S8.

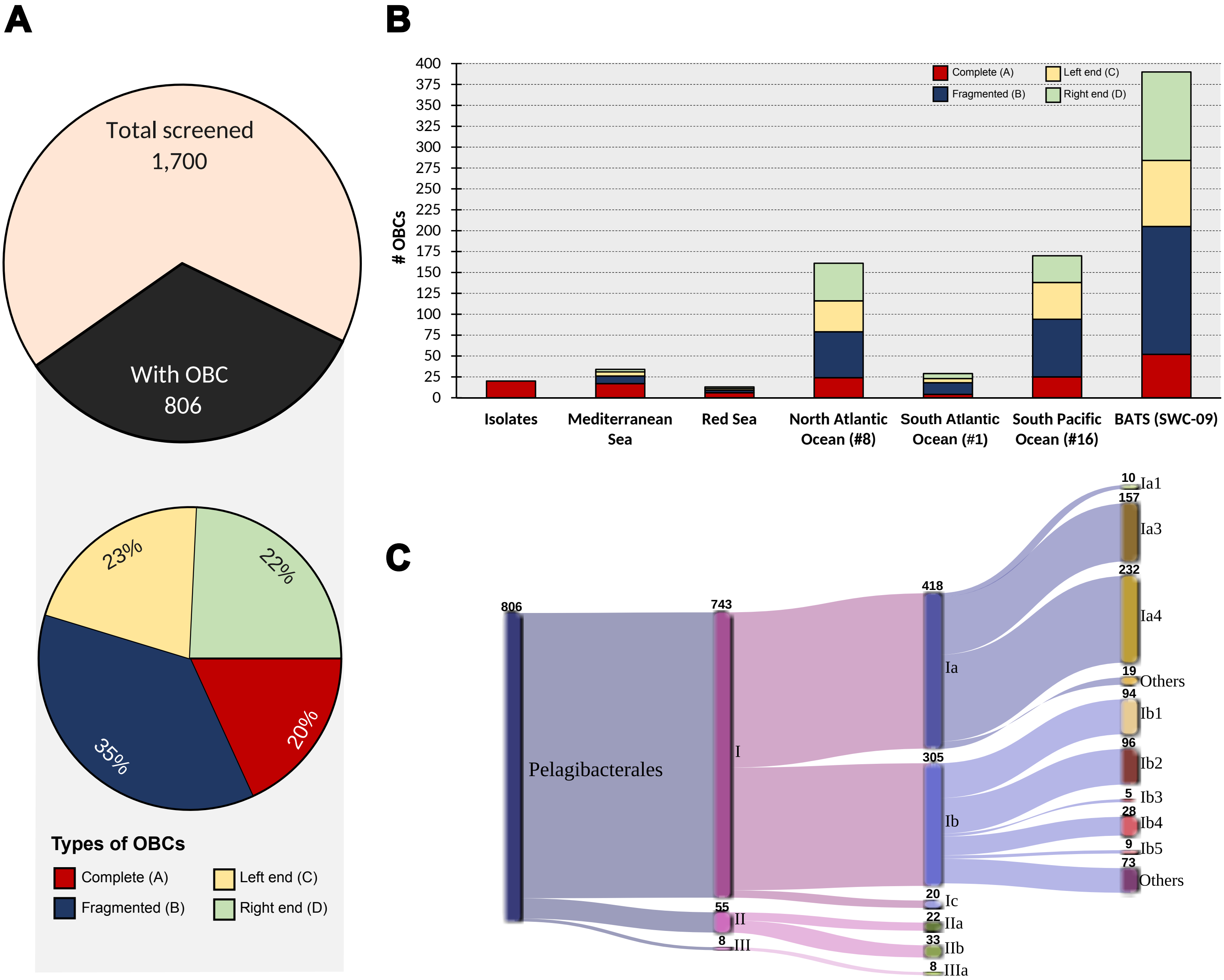


**Figure S1. A.** Upper pie chart indicates the total number of Pelagibacterales genomes screened (pale color), and the number of genomes on which we could identify the O-chain biosynthetic gene cluster (OBC) (black area). The bottom pie chart distributes the 806 OBCs according to their completeness: A – complete OBC; B – the boundaries of the OBC, i.e. the 23S rRNA gene on the left-hand side and the 5S rRNA gene on the right-hand side, were detected, but in two different contigs from the same genome; C – only the left-hand side; D – only the right-hand side. **B.** Number of OBCs recovered by oceanic region. **C.** Taxonomic classification of Pelagibacterales-containing OBCs based on a maximum-likelihood phylogenetic tree from shared proteins (see methods). The resulting phylogenetic groups follow the nomenclature described in Haro-Moreno et. al., 2020 (32).

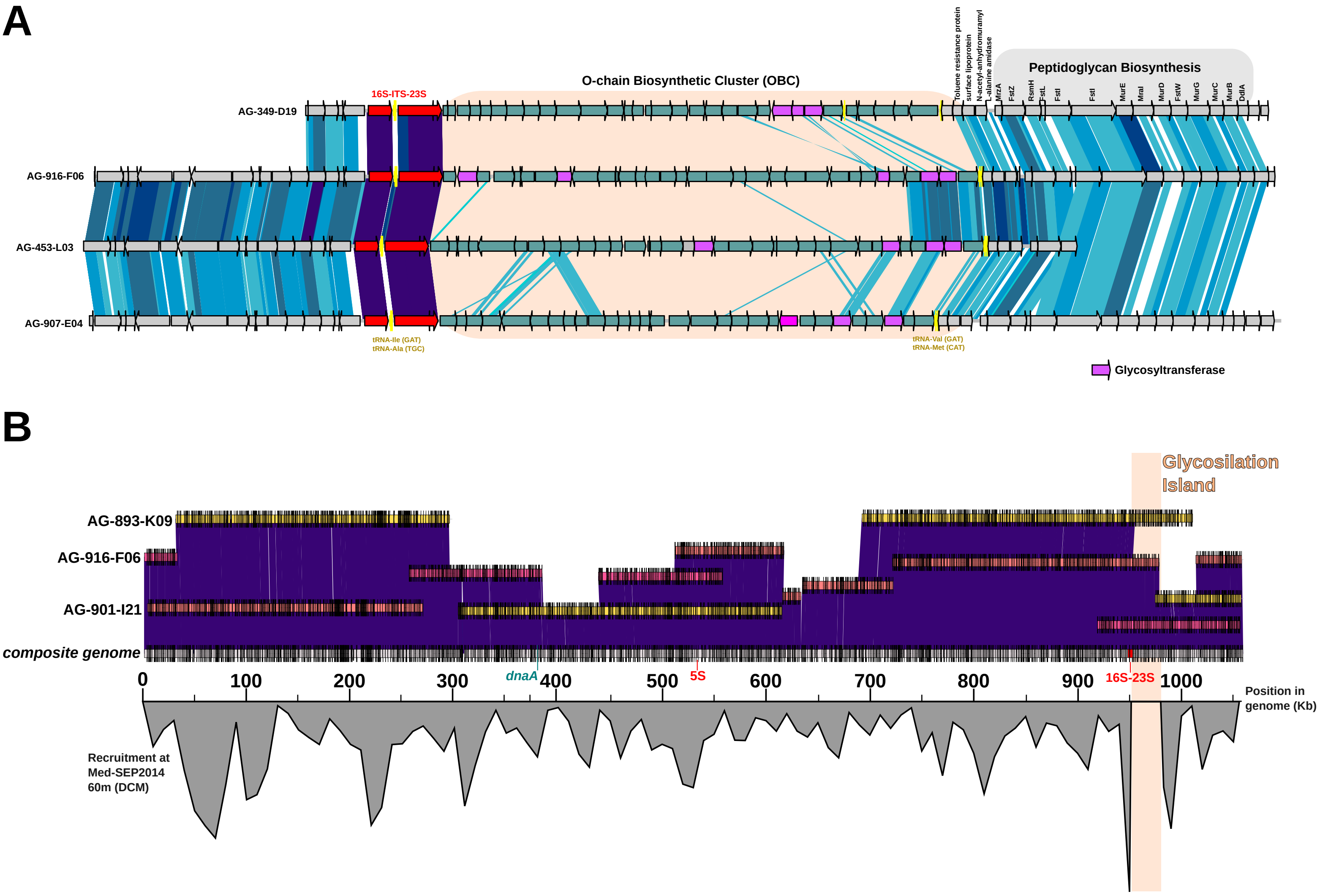


**Figure S2. A.** Genomic comparison of four selected complete OBCs from members of the Ib.4 phylogroup. Note that in this case, the glycosylation island is located between the 16S-ITS-23S rRNA genes and the tRNA-Val, tRNA-Met genes, and the core genes involved in the peptidoglycan biosynthesis. **B.** Reconstruction of a partial genome, Ib4-rB, in a single contig after the co-assembly of 3 nearly identical (>99 % ANI) SAGs. The locations of the dnaA, 16S, 23S, and 5S rRNA genes are indicated. The metagenomic fragment recruitment of this genome in the Mediterranean Sea (Med-SEP2014-60m) confirmed the location of the glycosylation island.

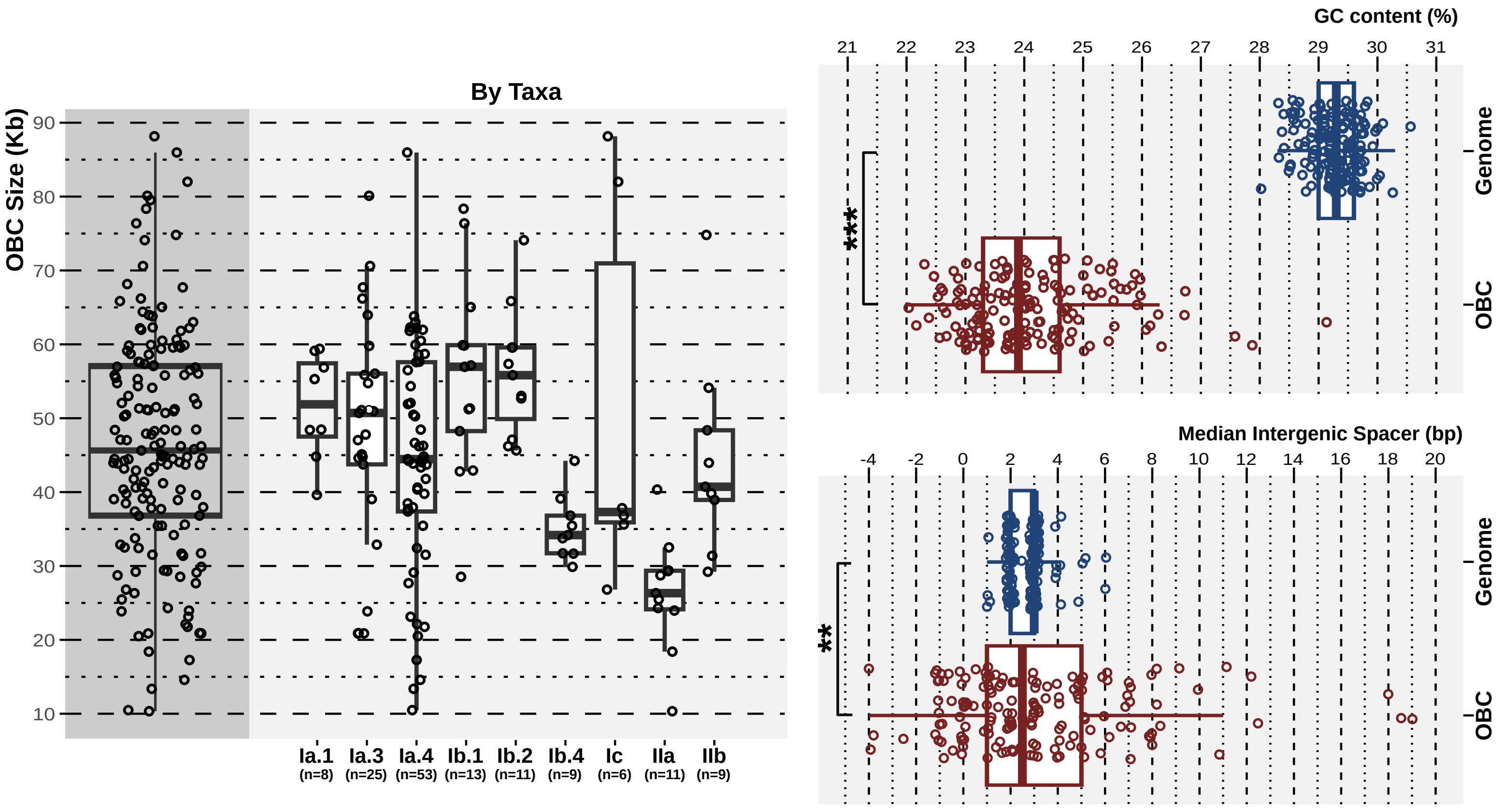


**Figure S3.** Genomic properties of the 163 complete OBCs and their corresponding genomes. Boxplot on the left summarises the median, first and third quartiles of the length of the OBC, considering all sequences as a unit (leftmost boxplot) or divided by subgroups. Numbers below the taxonomic name indicate the number of OBCs for each group. Boxplots on the right show the difference in the GC content and the intergenic spacer (upper and lower panels, respectively) between the OBC and its genome. Stars indicate the p-value (*** p-value < 0.001, ** p-value < 0.01).

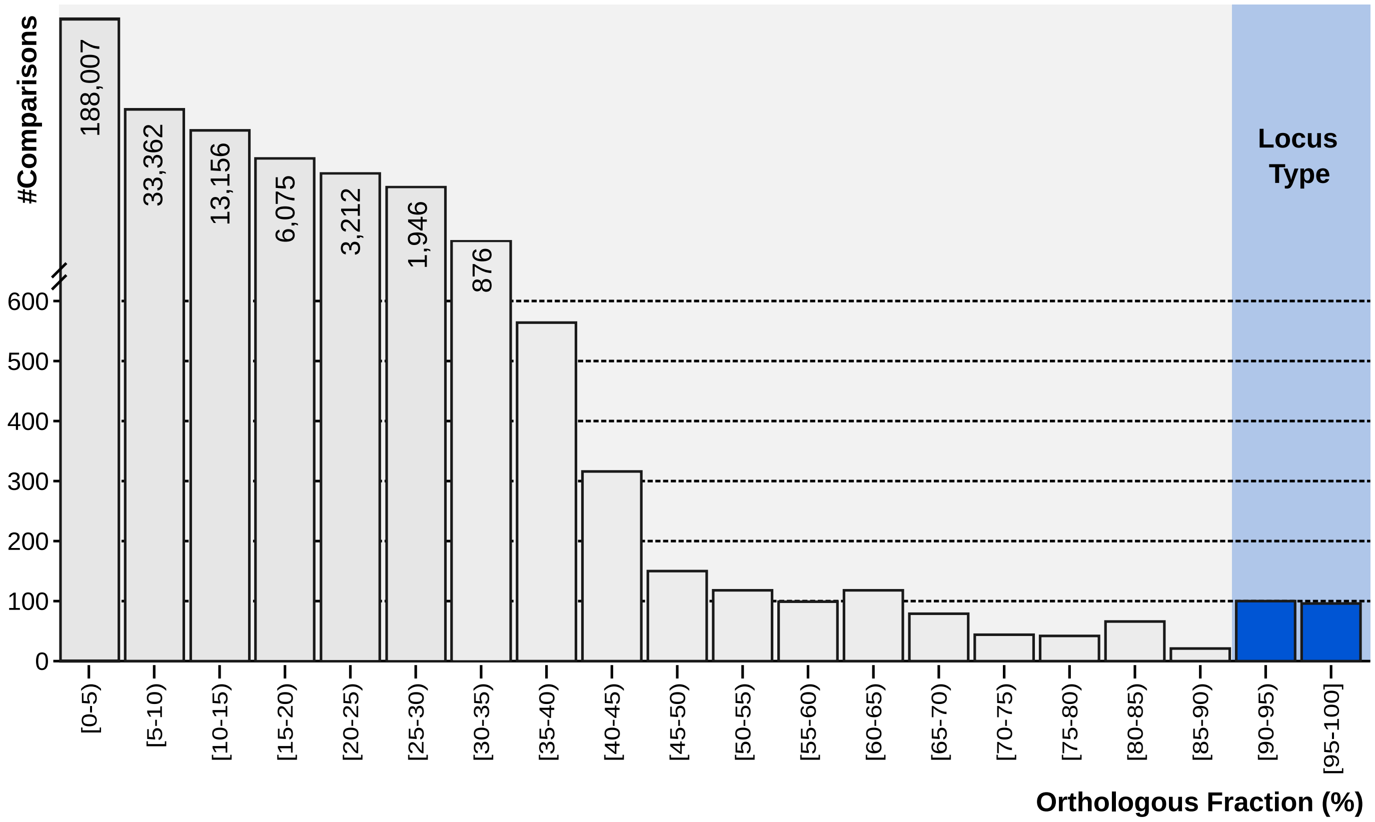


**Figure S4.** Histogram resulted from the all-vs-all comparison of the 806 OBCs at 50 % amino acid identity threshold. X-axis indicates the percentage of orthologous genes (OF) of the shortest sequence (genes shared between two OBCs), in groups of 5 % OF.

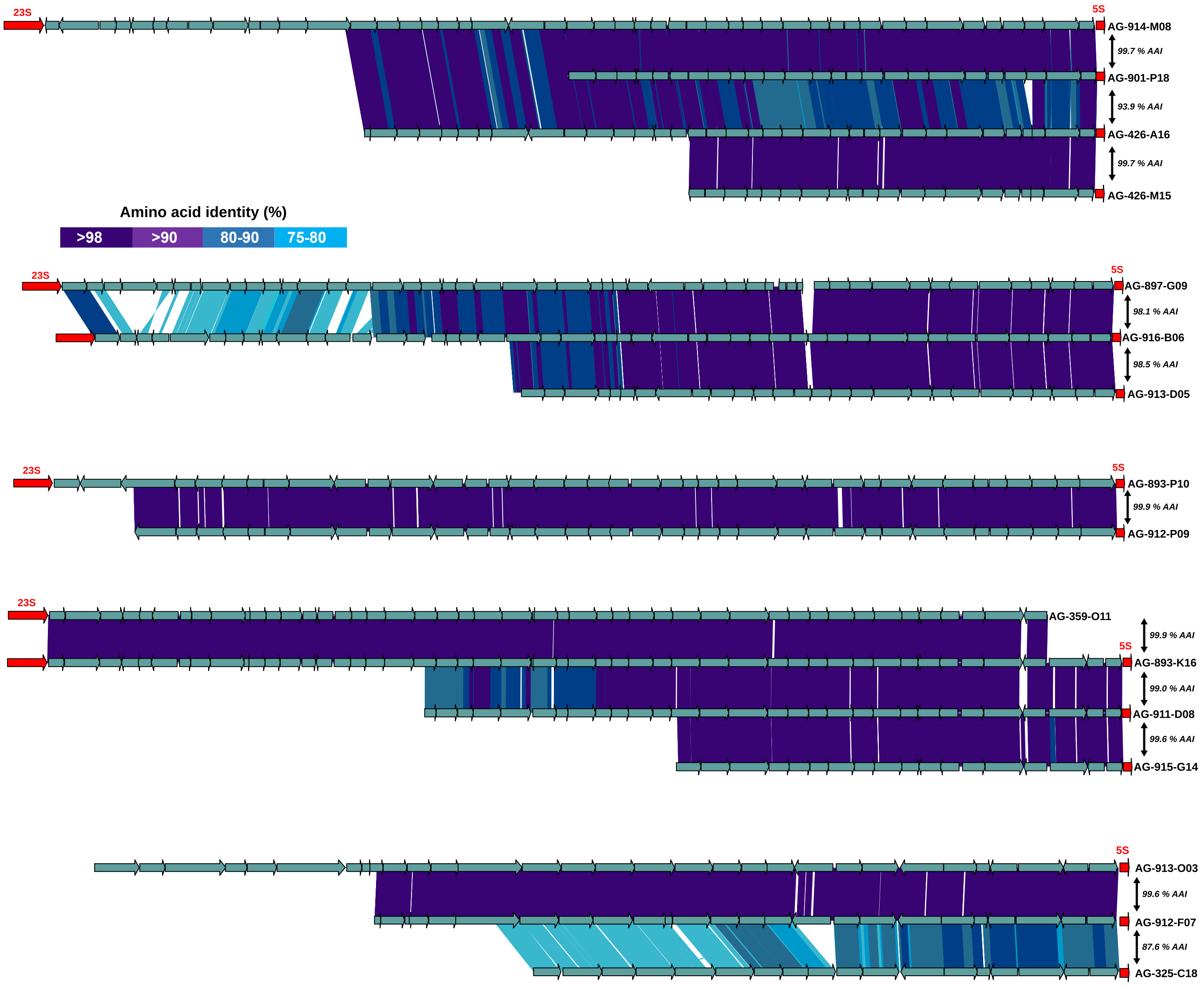


**Figure S5.** Examples of OBC-type sharing among Pelagibacterales genomes. OBCs are aligned and AAI color-coded.

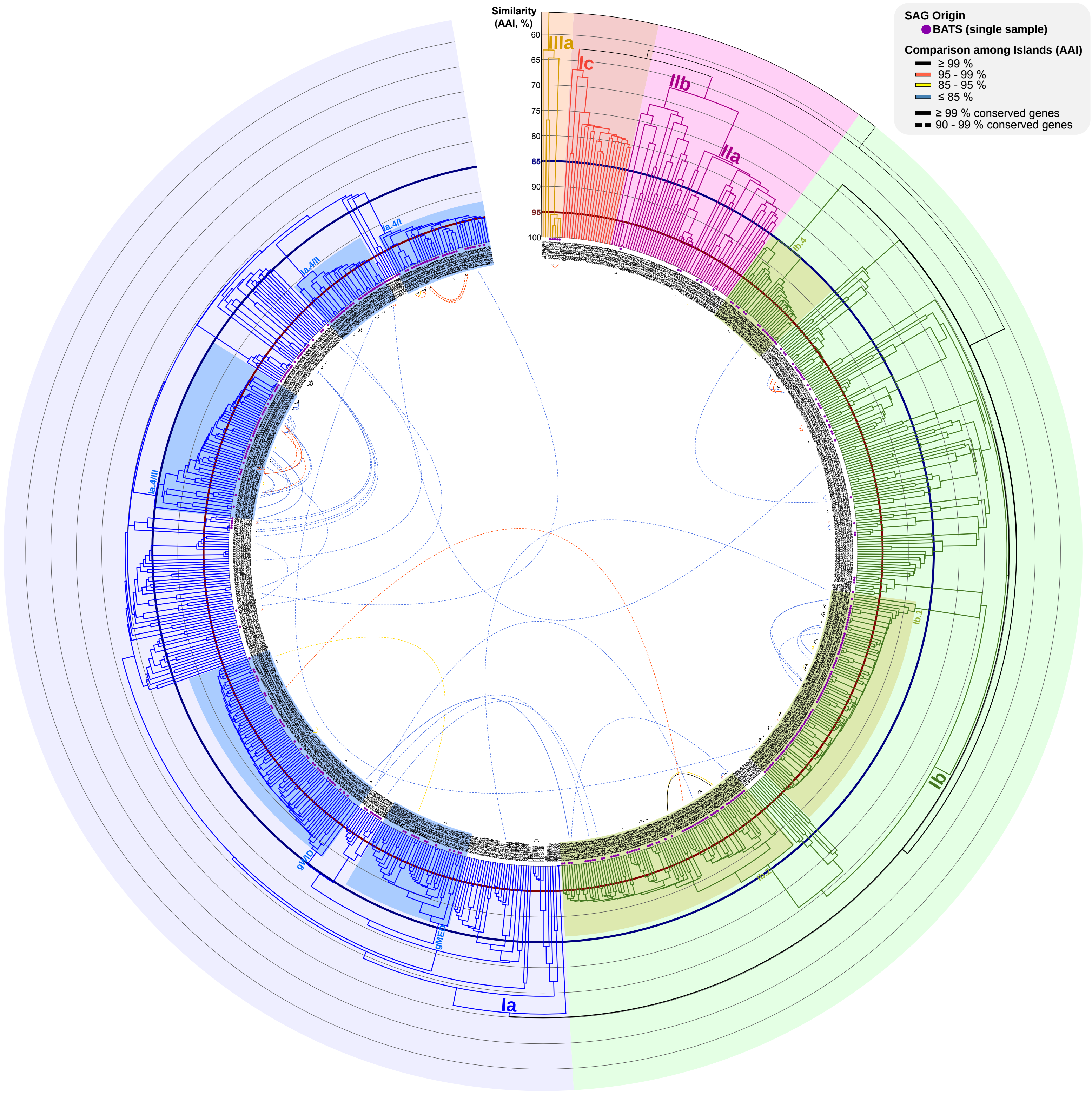


**Figure S6.** Cladogram-based classification of the 806 genomes containing an OBC, represented as in Figure 1.

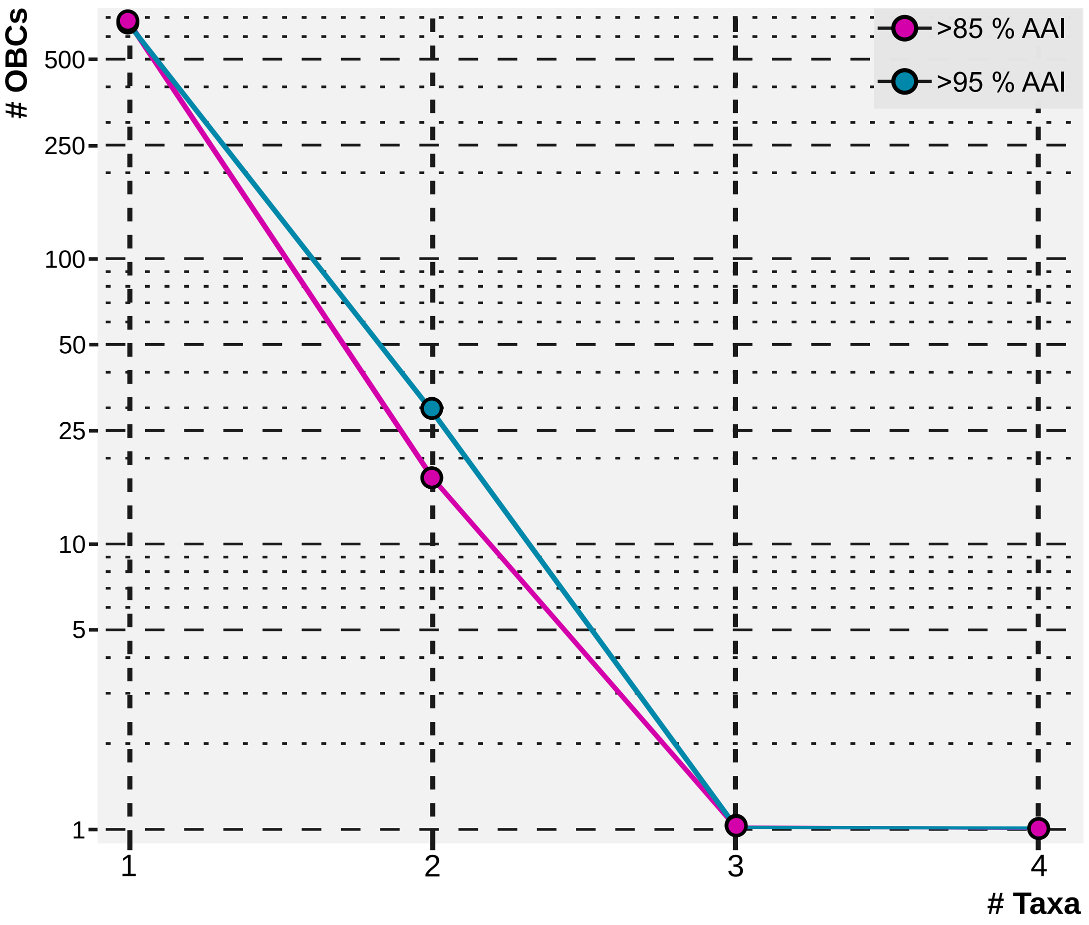


**Figure S7.** Number of OBCs found in one or several taxonomic groups (>85 % AAI or >95% AAI).

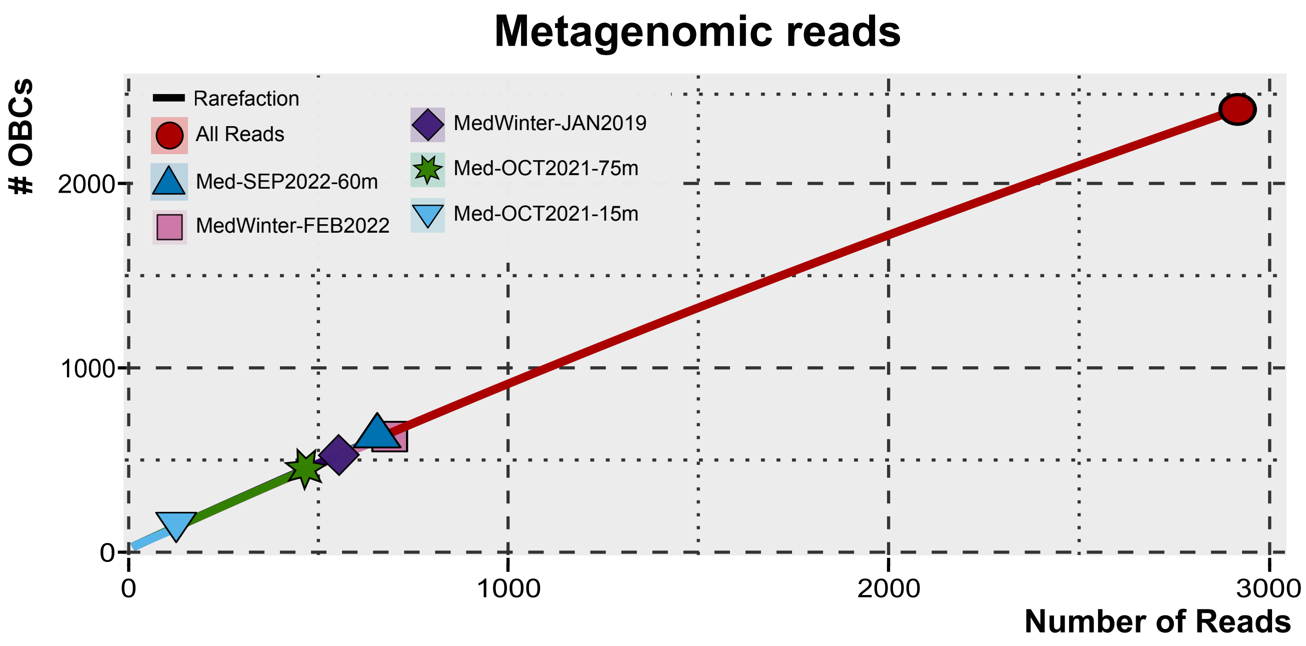


**Figure S8.** Rarefaction (solid line) curves based on OBC diversity against the number of sequences from a set of five PacBio Sequel II metagenomic reads collected from the Mediterranean Sea.
